# A Hyperactive Kunjin Virus NS3 Helicase Mutant Demonstrates Increased Dissemination and Mortality in Mosquitoes

**DOI:** 10.1101/2020.05.26.117580

**Authors:** Kelly E. Du Pont, Nicole R. Sexton, Martin McCullagh, Gregory D. Ebel, Brian J. Geiss

**Author notes:** Address correspondence to Brian J. Geiss,. K.E.D. and N.R.S. contributed equally to this work.

## Abstract

The unwinding of double-stranded RNA intermediates is critical for replication and packaging of flavivirus RNA genomes. This unwinding activity is achieved by the ATP-dependent nonstructural protein 3 (NS3) helicase. In previous studies, we investigated the mechanism of energy transduction between the ATP and RNA binding pockets using molecular dynamics simulations and enzymatic characterization. Our data corroborated the hypothesis that Motif V is a communication hub for this energy transduction. More specifically, mutations T407A and S411A in Motif V exhibit a hyperactive helicase phenotype leading to the regulation of translocation and unwinding during replication. However, the effect of these mutations on viral infection in cell culture and *in vivo* is not well understood. Here, we investigated the role of Motif V in viral replication using T407A and S411A West Nile virus (Kunjin subtype) mutants in cell culture and *in vivo.* We were able to recover S411A Kunjin but unable to recover T407A Kunjin. Our results indicated that S411A Kunjin decreased viral infection, and increased cytopathogenicity in cell culture as compared to WT Kunjin. Similarly, decreased infection rates in surviving S411A-infected *Culex quinquefasciatus* mosquitoes were observed, but S411A Kunjin infection resulted in increased mortality compared to WT Kunjin. Additionally, S411A Kunjin increased viral dissemination and saliva positivity rates in surviving mosquitoes compared to WT Kunjin. These data suggest that S411A Kunjin increases pathogenesis in mosquitoes. Overall, these data indicate that NS3 Motif V may play a role in the pathogenesis, dissemination, and transmission efficiency of Kunjin virus.

**IMPORTANCE:** Kunjin and West Nile viruses belong to the arthropod-borne flaviviruses, which can result in severe symptoms including encephalitis, meningitis, and death. Flaviviruses have expanded into new populations and emerged as novel pathogens repeatedly in recent years demonstrating they remain a global threat. Currently, there are no approved anti-viral therapeutics against either Kunjin or West Nile viruses. Thus, there is a pressing need for understanding the pathogenesis of these viruses in humans. In this study, we investigate the role of the Kunjin virus helicase on infection in cell culture and *in vivo*. This work provides new insight into how flaviviruses control pathogenesis and mosquito transmission through the nonstructural protein 3 helicase.

## INTRODUCTION

Kunjin virus, a West Nile virus (WNV) subtype, causes encephalitis epidemics in horses that are localized to Australia (1–4). Whereas, WNV has a much larger global impact present in almost every major continent except for South America and Antarctica (4, 5) and regularly results in encephalitis in humans as well as horses (6). Within the United States alone, approximately 3 million people are thought to have been infected with West Nile virus between 1999 and 2010 (7–9). Kunjin and WNV share a natural transmission cycle between *Culex* mosquito vectors and bird reservoir hosts (2). Humans and horses are considered dead-end hosts because they do not contribute to viral perpetuation. In humans, around 80% of WNV infected individuals are asymptomatic and the majority of symptomatic individuals experience a mild febrile illness. However, approximately 1:150 infections result in severe symptoms including meningitis and/or encephalitis, and ~9% of these cases are fatal (6, 10). Currently, there are vaccines against WNV for horses, but not for humans; no vaccines are available for Kunjin virus (5). Thus, there is a need for the development of vaccines and/or antiviral therapies for Kunjin and WNV infections. Developing a fundamental understanding of how Kunjin and WNV replicate within hosts, including the mosquito vector, is essential to the development of interventional strategies.

Kunjin and WNV belong to the *flavivirus* genus within the *Flaviviridae* family. *Flaviviridae* is a group of single-stranded positive-sense RNA viruses with genomes of approximately 11 kb in length (11–13). Kunjin virus is a subtype of WNV with a nucleotide and amino acid sequence identity of 82% and 93%, respectively (14–16). However, in humans, Kunjin virus results in low morbidity compared with WNV making it an excellent tool to study WNV replication with well-established molecular tools while minimizing risk (17). Additionally, Kunjin virus is less cytopathic than WNV, allowing for differences in virus-induced cell viability to be more easily visualized. Proteins and processes involved in viral replication are conserved across the flavivirus genus including for Kunjin, WNV, dengue, yellow fever, Japanese encephalitis, and Zika viruses (12, 18). Initially, the viral RNA genome is translated into a single polyprotein which is cleaved by host and viral proteases into three structural proteins (C, prM, and E) and eight nonstructural proteins (NS1, NS2A, NS2B, NS3, NS4A, 2K, NS4B, and NS5) (12, 18, 19). The viral NS replication proteins then generate a negative-sense anti-genomic RNA that is in complex with the positive-sense genomic RNA, forming the double-stranded RNA (dsRNA) intermediate complex (20, 21). The negative-sense anti-genomic RNA serves as a template for positive-strand synthesis (20); therefore, unwinding of the dsRNA intermediate is required for replication. Unwinding is achieved by the C-terminal helicase domain of NS3 (22–24).

NS3 helicase domain is a multi-functional viral protein that houses three enzymatic activities: RNA helicase, nucleoside triphosphatase (NTPase), and RNA 5’triphosphatase (RTPase) (25–28). NS3 helicase is a member of the superfamily 2 (SF2) helicases (29). The helicase domain consists of three subdomains (1, 2, and 3). Subdomains 1 and 2 are RecA-like structures that are highly conserved across all SF2 helicases, while subdomain 3 is unique to the viral/DEAH-like group of SF2 helicases (30). Additionally, there are eight structural motifs (Motifs I, Ia, II, III, IV, IVa, V, and VI) that are highly conserved across all viral/DEAH-like subfamilies with the SF2 helicases (29). These structural motifs are responsible for both substrate binding and enzymatic function within the helicase. The helicase domain is responsible for translocation and unwinding of the double-stranded RNA intermediate in an ATP-dependent manner during viral replication (31). Previous studies further identified Motif V as potentially critical for translocation and unwinding of the double-stranded RNA intermediate (32, 33). Motif V was described as a potential link between the ATP binding pocket and the RNA binding cleft through strong correlation between residues within Motif V and both binding pockets (32). The strongly correlated movements between ATP binding pocket and RNA binding cleft residues in our simulations suggest a physical linkage between the two sites that may be important for ATP driven helicase function. Additionally, mutants T407A and S411A in Motif V increased unwinding activity and decreased viral genome replication as compared to wild-type (WT), suggesting that the hydrogen bond between these two residues in WT inhibits helicase unwinding activity *in vitro* and *in vivo* (33). These data suggested that Motif V may serve as a molecular throttle on NS3 helicase function, but what effect these residues play on the larger viral replication cycle was not clear.

To better understand the effects NS3 Motif V mutations have on flavivirus replication, we sought to investigate the role of Motif V T407 and S411 residues on helicase function in cell culture and *in vivo* by introducing alanine mutations in full-length infectious Kunjin virus: T407A Kunjin and S411A Kunjin. Only the S411A Kunjin was recovered and it resulted in reduced viral yields compared with wild-type (WT) Kunjin. Additionally, S411A Kunjin showed increased cytopathic effect in comparison to WT Kunjin in cell culture. Similarly, when WT or S411A Kunjin viruses were intrathoracically injected into *Culex quinquefasciatus* mosquitoes, S411A Kunjin resulted in increased mortality compared with WT Kunjin. Upon further investigation of mosquito infection, S411A Kunjin viruses were found to disseminate and transmit more effectively than WT Kunjin viruses, even though the overall infection rate was lower than WT Kunjin. Overall, our data suggest that flaviviruses may use NS3 Motif V to help control cytotoxicity induced by NS3 during infection and limit virus-induced mortality in mosquito vectors.

## RESULTS

### S411A Kunjin virus increases cytopathic effect in cell culture

Previously, Motif V residues, T407 and S411, were mutated to alanine to disrupt a hydrogen bond that potentially stabilizes the Motif V secondary structure of NS3 helicase during viral replication (Fig. 1). These mutations were shown to decrease viral genome replication in a replicon-based system, while increasing helicase unwinding activity biochemically (33). In the present study, we introduced these mutations into the full-length infectious Kunjin virus to investigate the effects of these mutations on infectivity compared to WT Kunjin both in cell culture and in mosquito infections. We utilized a novel mutagenesis and a bacteria-free viral launch system to generate the T407A Kunjin and S411A Kunjin viruses in Vero cells. The first generation of S411A Kunjin was recovered from infection and the presence of the alanine mutation was verified with sequencing (Fig. 2). On the other hand, we were unable to recover the T407A Kunjin despite repeated attempts, which was consistent with our previously reported decrease in T407A viral genome replication in replicon assays (33). Second generation stocks of WT Kunjin and S411A Kunjin were generated and the viruses were titered for further experiments. We noted the plaque morphology for both WT Kunjin and S411A Kunjin (Fig. 3). WT Kunjin showed large, faint plaque sizes (Fig. 3A), while S411A Kunjin showed small, but distinctly clear plaques (Fig. 3B), suggesting a potential decrease in viral cell-to-cell spread and an increase in cytopathic effect for S411A Kunjin infected cells compared to WT Kunjin. Since these results suggest that S411A Kunjin may be more toxic to cells during infection, we further investigated the effect of the S411A Kunjin on cell viability.

**FIG. 1.**
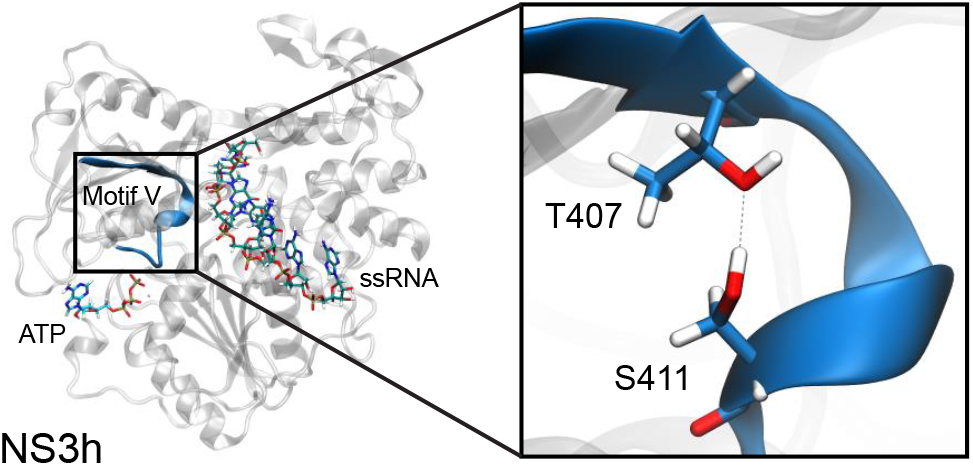
S411 and T407 interaction within Motif V in NS3 helicase. Residues T407 and S411 interact with each other through a hydrogen bond within Motif V.

**FIG. 2.**
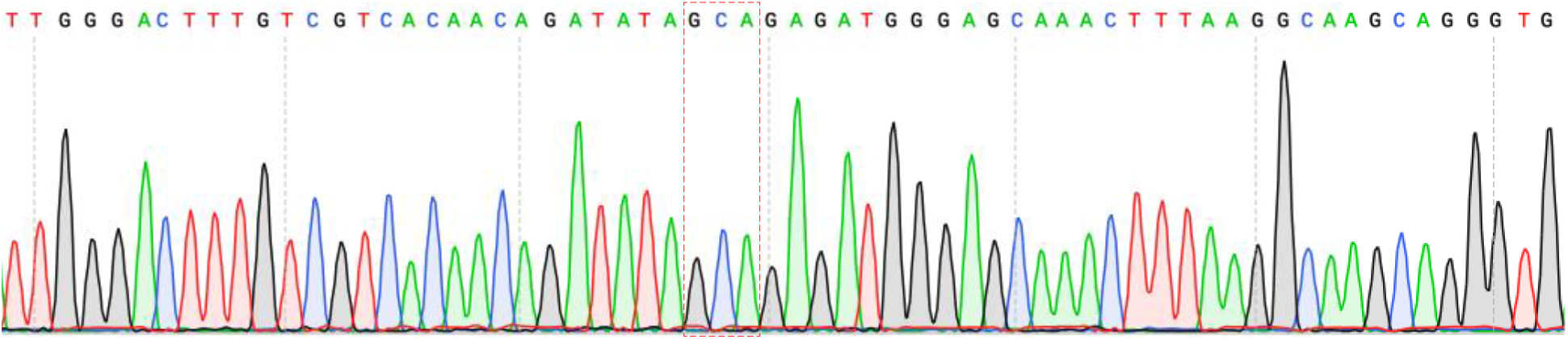
Verification of alanine mutation in S411A Kunjin virus via Sanger sequencing. Results from Sanger sequencing verifies alanine mutation for position 411 through the presence of the alanine codon (highlighted in red box). The original serine codon within the red box was TCT. Two nucleotides were changed to introduce the alanine mutation. Refer to GenBank accession number (<u>AY274504.1</u>) for wild-type Kunjin FLSDX.

**FIG. 3.**
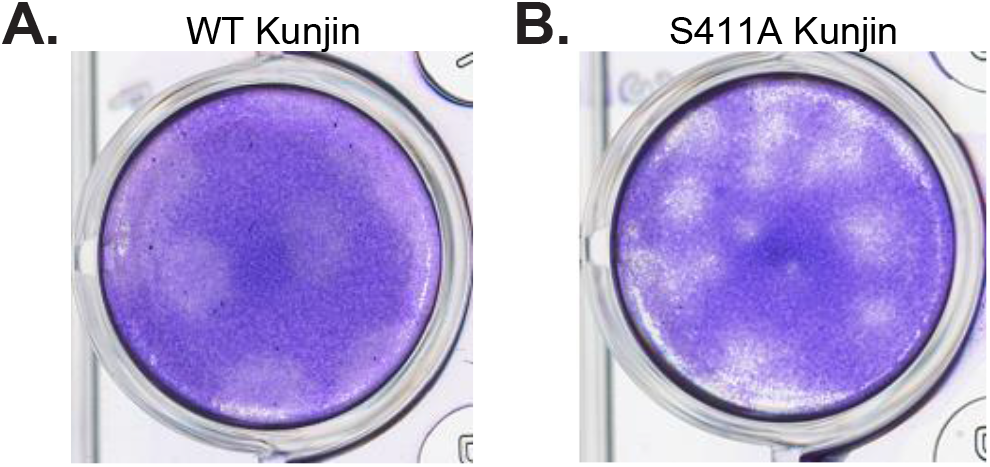
Plaque morphology suggests an increased cytopathic effect for S411A Kunjin. Viral titers were obtained for WT and S411A Kunjin viruses and the plaque morphology is shown for A) WT Kunjin and B) S411A Kunjin.

### S411A Kunjin reduces NADH and intracellular ATP levels leading to increased cellular death

We utilized resazurin and CellTiter-Glo assays to quantify virus-induced cell killing in HEK293T and Vero cells infected with either WT Kunjin or S411A Kunjin at a multiplicity of infection (MOI) of five PFU/cell. Both of these assays estimate cell viability through the measurement of metabolically active cells using fluorescence and luminescence, respectively. In the resazurin assay, resazurin, a nonfluorescent dye, converts to resorufin, a highly fluorescent dye, in response to the reducing environment of heathy, growing cells (34–36). We measured the relative fluorescence units (RFU) of resazurin in uninfected, WT Kunjin, or S411A Kunjin infected Vero and HEK293T cells every 24 hours for six days (Fig. 4A and B). We also measured media as a negative control to determine the baseline media fluorescence. The cell viability measurements of uninfected Vero and HEK293T cells increased gradually over the duration of the experiment suggesting that the cells are healthy and growing for the entirety of the experiment. The cell viability measurements during the first 72 hours for WT Kunjin infection in Vero and HEK293T cells were similar to that of uninfected cells. However, cell viability measurements were lower in fluorescent signal compared to uninfected cells. After 72 hours post infection (p.i.), cell viability measurements for WT Kunjin infections continued to increase in fluorescence reaching 7.5 ± 0.3 ×10^5^ RFU at 120 hours p.i. for Vero cells and 7.8 ± 0.2 ×10^5^ RFU at 96 hours p.i. for HEK293T cells. After which point, cell viability measurements decreased in fluorescence by 144 hours p.i. suggesting that WT Kunjin induced cell toxicity is overtaking cellular replication. In the case of S411A Kunjin infected Vero and HEK293T cells during the first 72 hours, cell viability measurements demonstrated similar levels of fluorescence to that of uninfected cells. Although the cell viability measured for S411A Kunjin was decreased compared to uninfected cells. As the S411A Kunjin infection continued, cell viability measurements significantly reduced in fluorescence between 96 and 144 hours p.i. ending with 5.3 ± 0.3 ×10^5^ RFU for Vero cells and 5.7 ± 0.2 ×10^5^ RFU for HEK293T cells. Together, these data suggest that cells are relatively healthy in Kunjin infected cells for at least the first 72 hours in Vero and HEK293T cells; after which point population cell viability in S411A Kunjin infected cells is negatively affected immediately in both cell lines, whereas a 24 hour and 48 hour delay are observed for decreased cell viability measurements with WT Kunjin infection for HEK293T and Vero cells, respectively.

**FIG. 4.**
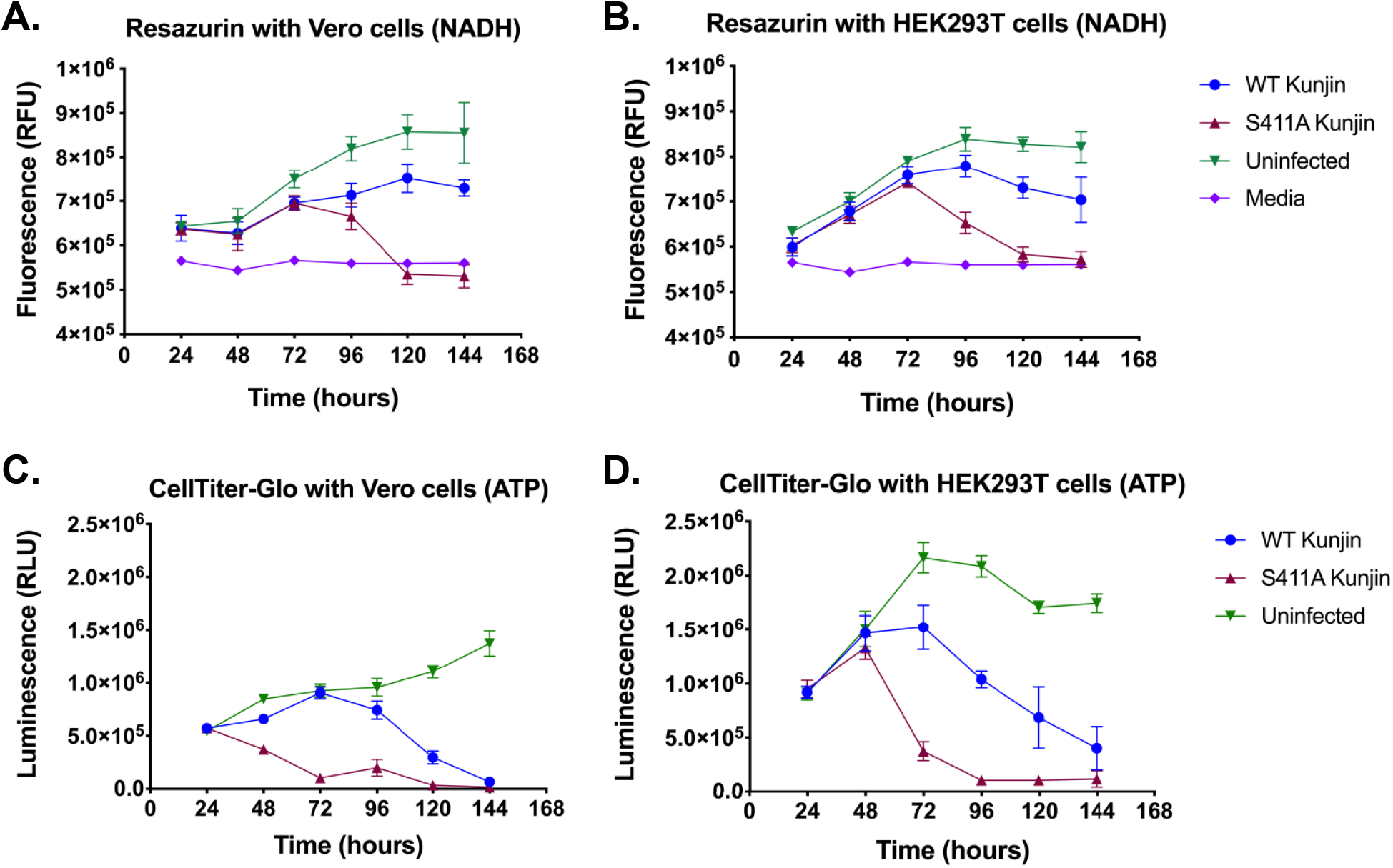
S411A Kunjin decreases cell viability. WT Kunjin and S411A Kunjin infected A) Vero cells and B) HEK293T cells were measured for cellular metabolism through resazurin. Similarly, WT Kunjin and S411A Kunjin infected C) Vero cells and D) HEK293T were measured for intracellular ATP levels through CellTiter-Glo. All infections were performed at a MOI of five PFU/cell.

Another way to infer metabolically active cells or cell viability is through detection of intracellular ATP levels. We utilized the CellTiter-Glo assay which uses the luciferase reaction, an ATP-dependent reaction, to convert luciferin to oxyluciferin and several byproducts including light (34). The byproduct, light, was measured in relative luminescence units (RLU) for uninfected, WT Kunjin or S411A Kunjin infected Vero and HEK293T cells every 24 hours for six days (Fig. 4C and D). Over the course of the experiment, uninfected Vero cells progressively increased in luminescence from 5.5 ± 0.3 ×10^5^ to 1.4 ± 0.1 ×10^6^ RLU (Fig. 4C) suggesting that the uninfected cells were healthy and metabolically active for the six-day experiment. However, cell viability measurements of uninfected HEK293T cells increased linearly for the first 72 hours; after which point, the cell viability measurements decreased and then leveled off at 1.7 ± 0.07 ×10^6^ RLU (Fig. 4D), suggesting that uninfected HEK293T cells become less metabolically active after 96 hours compared to the Vero cells. As for infection with WT Kunjin, the cell viability measurements steadily increased for the first 72 hours for Vero cells and for the first 48 hours for HEK293T cells similar to the observed cell viability measurements of uninfected Vero and HEK293T cells. At 96 hours p.i. in Vero cells and 72 hours p.i. in HEK293T cells, cell viability measurements of WT Kunjin infected cells decreased compared to uninfected cells. The population cell viability of WT Kunjin infected cells continued to decrease reaching 6.5 ± 3.0 ×10^4^ RLU in Vero cells and 4.0 ± 2.0 ×10^5^ RLU in HEK293T cells at 144 hours. These data suggested that infection with WT Kunjin negatively affected cell viability after 72 hours p.i. compared to uninfected cell viability. On the other hand, cell viability measurements with S411A Kunjin infection decreased after 24 hours p.i. in Vero cells and after 48 hours p.i. for HEK293T cells. For the remainder of the experiment, the population cell viability continued to decrease in S411A Kunjin infected Vero and HEK293T cells suggesting that both Vero and HEK293T cells are extremely sensitive to S411A Kunjin and thus cell viability is significantly reduced in the presence of the mutated virus. Together, these results suggest that infection with S411A Kunjin in either Vero or HEK293T cells negatively affected cell viability more quickly than infection with WT Kunjin.

### S411A Kunjin results in decreased and delayed viral replication kinetics

The results presented in the previous section indicated that S411A Kunjin induced increased cellular death during infection. This prompted the question: how does increased cellular death resulting from infection with S411A Kunjin affect replication kinetics of the virus? Therefore, we performed a multi-step replication kinetics experiment with WT or S411A Kunjin infected HEK293T cells at a MOI of 0.01 PFU/cell over a five day period. Every 12 hours viruses were collected and viral titers were determined via focus forming assays (Fig. 5). At 12 hours post infection, the WT and S411A Kunjin viral titers were not significantly different. At 24 hours p.i., S411A Kunjin remained in the lag phase while WT Kunjin had entered the exponential replication phase, demonstrating delayed replication with the S411A Kunjin infection. Over the last four days of infection, S411A Kunjin maintained and expanded the initial delay in exponential replication and reached an ~1 log lower peak viral titer compared to WT Kunjin. Overall, these data suggest that S411A Kunjin does not replicate as efficiently as WT Kunjin. These results are consistent with data reported by Du Pont *et al.*, suggesting that the increased helicase unwinding activity seen with the recombinant S411A NS3 helicase negatively affects viral replication in fully infectious S411A Kunjin virus (33). Considering the observations that S411A Kunjin resulted in decreased viral replication and increased cellular death, we next investigated the effects of the S411A mutation on Kunjin infection *in vivo*.

**FIG. 5.**
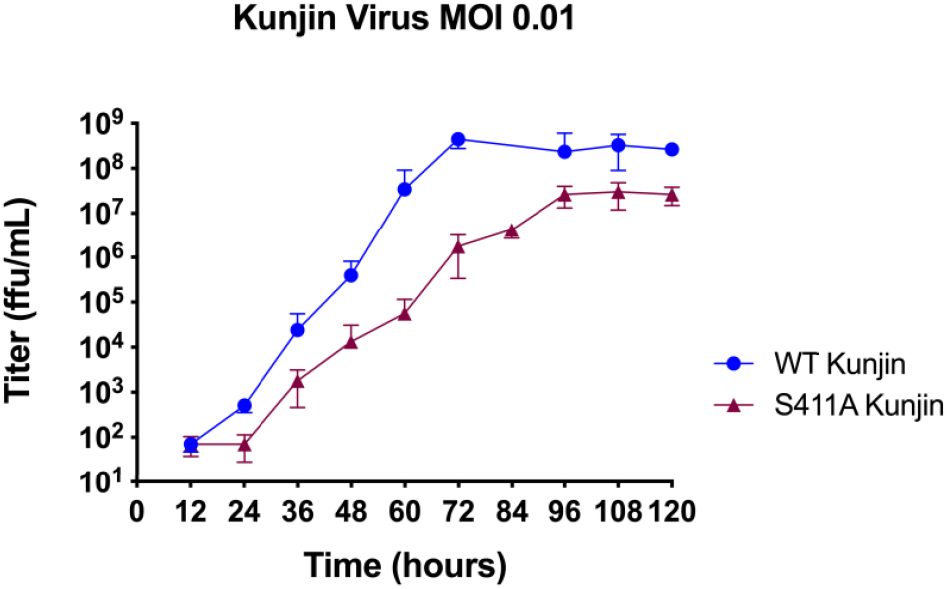
S411A Kunjin decreases and delays viral replication kinetics. Replication kinetics experiments were performed for WT and S411A Kunjin viruses. HEK293T cells were infected at a MOI of 0.01 PFU/cell.

### S411A Kunjin results in increased mortality in mosquitoes compared to WT Kunjin when IT injected but not when bloodfed

For the *in vivo* studies, we did not have access to a colony of *Cx. annulirostris* mosquitoes, the primary vector for Kunjin virus, but we had an established colony of *Cx. quinquefasciatus* that are infectable by Kunjin virus. *Cx. quinquefasciatus* mosquitoes were bloodfed with defibrinated calf’s blood diluted by half with titer equilibrated WT Kunjin, S411A Kunjin, or media alone as a negative control. Similarly, female *Cx. quinquefasciatus* mosquitoes were subjected to intrathoracic injection (IT) of 345 plaque forming units (PFU) per mosquito of WT Kunjin, S411A Kunjin, or conditioned media. Mosquito mortality was recorded daily for 15 or 9 days, respectively. Overall, virus exposed mosquito mortality was low in both the bloodfed and IT injected cohorts (Fig. 6), consistent with previous observations of Kunjin virus in *Cx. quinquefasciatus* mosquitoes (37). When bloodfed, no difference was observed in mortality rates for mosquitoes exposed to WT Kunjin vs. S411A Kunjin. However, the small rate of mortality for virus exposed mosquitoes (~10%) was significantly different from mosquitoes exposed to media alone (Fig. 6A). In contrast with bloodfed data but consistent with cell culture and replication kinetics data, when virus was introduced through IT injection, to bypass the midgut barrier, only S411A Kunjin resulted in increased mortality (Fig. 6B). Together these data suggest that S411A Kunjin is more lethal to mosquitoes than WT Kunjin once the virus has been able to establish infections and/or transverse through the mosquito midgut barrier. This result led us to further investigate the specifics of infection of *Cx. quinquefasciatus* by WT and S411A Kunjin viruses.

**FIG. 6.**
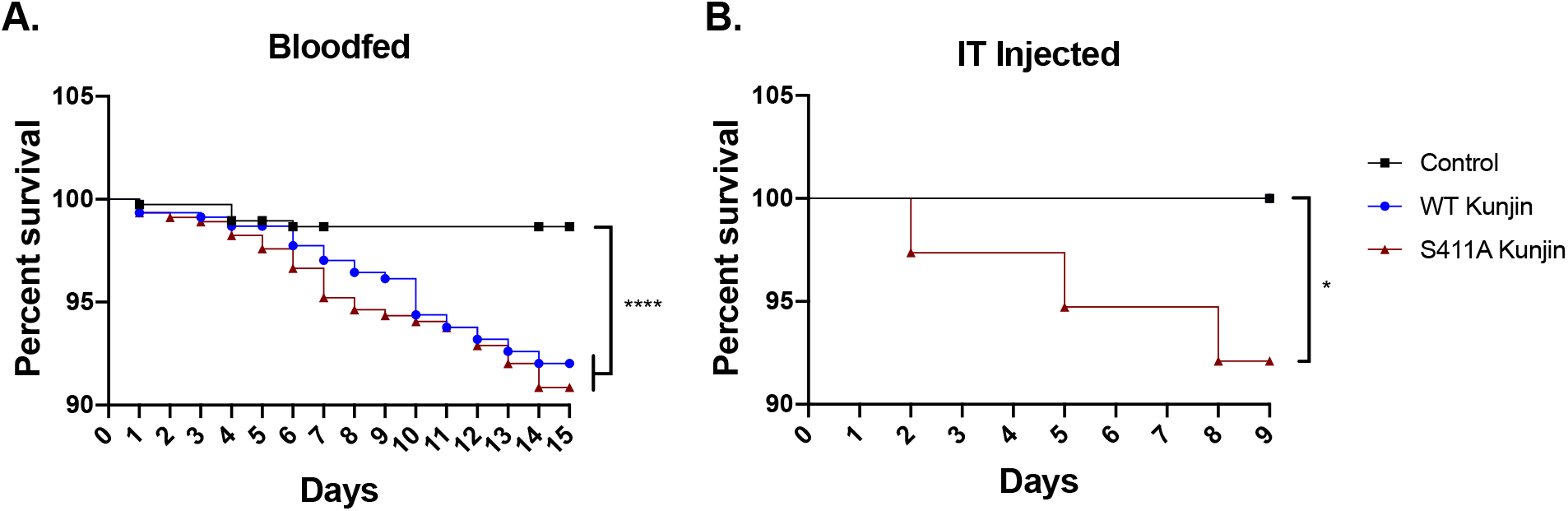
S411A Kunjin viruses are more lethal to *Cx. quinquefaciatus* mosquitoes than WT Kunjin. Female *Cx. quinquefaciatus* mosquitoes were exposed to WT (blue circles) or S411A (red triangles) Kunjin virus through either A) infectious bloodmeals, or A) by IT injection. Control mosquitoes were exposed to bloodmeals containing media or injected with media alone. Mortality was recorded daily for 15 or 9 days respectively. Survival curves compared by Logrank test for trend (P<0.0001 = ****, P<0.05 = *) A) n = 425/condition, B) n = 40/condition.

### S411A Kunjin has a lower infection rate but disseminates more efficiently than WT Kunjin

Similar to the mortality experiments, *Cx. quinquefasciatus* mosquitoes were infected with either WT Kunjin or S411A Kunjin by bloodmeal. Mosquito legs/wings, saliva, and bodies were collected after 7 days and determined to be positive or negative for infection by plaque assay. While ~58% of mosquitoes infected with WT Kunjin were positive for the virus at day 7, only ~8% of mosquitoes infected with S411A Kunjin were positive (Fig. 7A). Dissemination was inefficient for WT Kunjin with only 6% of mosquitoes having positive titers in the legs and wings, demonstrating a strong barrier to escape from the midgut. Similarly, less than 2% of infected mosquitoes resulted in positive saliva samples (Fig. 7A). Despite low infection rates for mosquitoes infected with S411A Kunjin, positive legs/wings and saliva were identified across multiple replicate experiments, with nearly 50% of infected mosquitoes having disseminated virus and 50% of those with disseminated virus having positive saliva. These data led to the question: does S411A Kunjin allow for higher relative rates of dissemination?

**FIG. 7.**
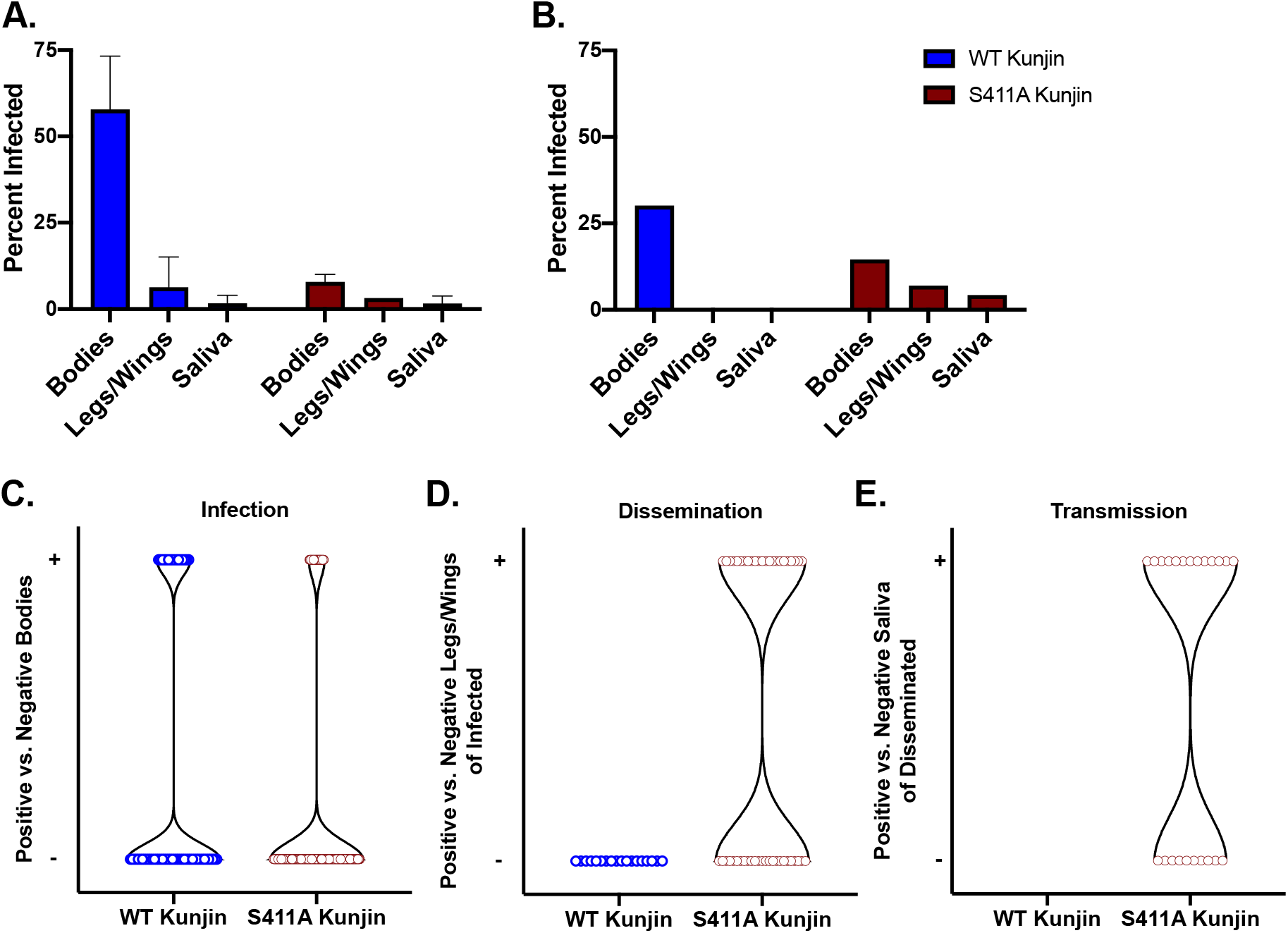
The S411A Kunjin is less capable than WT Kunjin of infecting mosquitoes but disseminates and transmits more efficiently once established. Engorged female *Cx. quinquefascitus* mosquitoes exposed to infectious bloodmeals containing either WT or S411A Kunjin virus were housed for A) 7 or B-E) 14 days post bloodfeed. Mosquitoes were dissected and legs/wings, saliva and bodies were collected and tested for the presence of Kunjin virus by plaque assay. Data is shown as A and B) percent of total exposed infected, C) total negative and positive bodies, D) positive legs/wings from total infected, or E) total positive saliva from total disseminated. A) n = 64/condition, B) WT Kunjin n = 60, S411A Kunjin n = 390.

To answer this question a second, much larger cohort of *Cx. quinquefasciatus* mosquitoes were infected by bloodmeal with WT Kunjin or S411A Kunjin. Enough mosquitoes were dissected to generate and estimated 30 infected mosquitoes per condition: 60 exposed to WT Kunjin and 390 exposed to S411A Kunjin. Since mosquitoes continue to die up to 14 days post bloodfeed, mosquitoes were collected at 14 days post blood meal instead of 7 days in an attempt to assure sufficient numbers of S411A Kunjin infected mosquitoes. Again, WT Kunjin was observed to infect a larger percent of exposed mosquitoes compared with S411A Kunjin (~30% vs. ~15%) (Fig. 7B,C), whereas, S411A Kunjin demonstrated higher rates of dissemination compared with WT Kunjin (Fig. 7B,D). No legs/wings or saliva samples from WT Kunjin infected mosquitoes were found to be positive at 14 days post blood meal (Fig. 7B,D,E). In contrast and supporting these data from smaller cohorts collected at 7 days post blood meal, 48% of S411A Kunjin infected mosquitoes had infected legs/wings and 61% of mosquitoes with S411A Kunjin infected legs/wings resulted in positive saliva samples. These data demonstrate that the S411A Kunjin was less capable of infecting *Cx. quinquefasciatus* via blood meal compared with WT Kunjin. However, these data also suggest that when S411A Kunjin was able to establish infection in *Cx. Quinquefasciatus* mosquitoes it is able to escape the midgut barrier more efficiently than WT Kunjin, resulting in dissemination, infection of the salivary glands, and delivery to the saliva. Finally, when considered in combination with the survival data, these data further support that when S411A Kunjin was able to establish infection in *Cx. quinquefasciatus* mosquitoes it is more lethal.

## DISCUSSION

Previous work by our group has supported the hypothesis that Motif V in flavivirus NS3 helicase is a communication hub for translocation and unwinding of the dsRNA intermediate during flavivirus replication (32, 33). More specifically, we found that NS3 Motif V residues T407 and S411 exhibit an increased helicase unwinding activity in biochemical assays when mutated to alanine residues, while we observed a reduction in replication of T407 and S411 mutant replicons. These previous results suggest that T407 and S411 are responsible for regulating NS3 helicase function during flavivirus replication. In this study we further investigated the role of T407 and S411 helicase residues in the full-length infectious Kunjin virus in cell culture and *in vivo* experiments. S411A Kunjin was successfully recovered and confirmed via sequencing (Fig. 2). However, T407A Kunjin was not recovered which was consistent with the previous results indicating ablated viral genome replication activity (33). We utilized WT Kunjin and S411A Kunjin in several cell culture experiments including viral replication, resazurin and CellTiter-Glo assays. Additionally, we compared WT Kunjin and S411A Kunjin in several *in vivo* experiments including infection, dissemination and transmission within *Cx. quinquefasciatus* mosquitoes. We observed that the S411A Kunjin reduced cell viability during infection leading to increased cytopathic effect observed in the plaque morphology and several metabolic assays in cell culture. Additionally, results demonstrated a lower initial infection rate for S411A Kunjin within mosquitoes but once infection is established efficient dissemination occurs compared with WT Kunjin infections, potentially causing the observed increased mortality rates in mosquitoes. Overall, our data suggest that the NS3 S411 in Motif V influences infection induced cellular death and subsequent mortality in mosquito vectors.

Plaque morphology of viruses is a classical indicator of the effects of a mutation on viral cytopathic effect in cells and spread between cells. We observed large and fuzzy plaques with WT Kunjin, while S411A Kunjin plaques were small and clearly defined (Fig. 3), suggesting that S411A Kunjin is more toxic to cells, but is not able to spread as rapidly as WT Kunjin. Our previous work had indicated that the S411A mutation in a replicon-based system reduced viral genome replication (33), so the small plaque size was expected. However, the formation of clearer plaques was not. Therefore, we performed a more quantitative investigation of S411A Kunjin effect on cell viability using two assays (resazurin and CellTiter-Glo) that probed for different aspects of metabolically active cells, NADH content and ATP content. The results from both assays indicated that infection with S411A Kunjin results in a larger decrease in metabolic activity compared to WT Kunjin within both HEK293T and Vero cells (Fig. 4). Previously, studies have shown that reduced intracellular ATP levels leads to proteasome inhibition that induces apoptosis leading to cellular death (38–43). Therefore, our metabolic activity data is consistent with our plaque morphology data in that infection with S411A Kunjin results reduced intracellular ATP levels and increased cytopathic effect through increased cell death. S411A Kunjin exhibited delayed and decreased viral replication kinetics compared to WT Kunjin (Fig. 5) suggesting that even though the mutated Kunjin virus is more toxic to cells, it does not replicate as efficiently as WT Kunjin. These data are consistent with previous studies reporting a decrease in viral genome replication with S411A helicase replicon (33).

An interesting but different hypothesis is that hyperactive NS3 helicase affects cellular mRNA. Studies on NS3 helicase function have focused primarily on its effect on genome replication and packaging (44), but our finding that a NS3 hyperactive helicase mutant increases cell death opens up the possibility that NS3 has roles in altering cellular physiology as well. Previously observed results indicated that recombinant NS3 S411A helicase mutant had a higher helicase rate but did not have a significantly higher ATPase rate (33), so it is unlikely that reduction of cell viability was due to decreased ATP from NS3 ATP degradation. However, it is possible that increased cytotoxicity is due to another effect of helicase activity on cellular physiology. The hyperactive NS3 helicase may be interacting with cellular RNAs leading to dysregulation of cellular homeostasis. NS3 could bind to cellular mRNAs and unwind their secondary structures, causing a disruption in RNA stability and recruitment of translational factors. This unwinding of cellular mRNAs would result in an imbalance within the cell inducing cellular apoptosis. We are currently exploring if NS3 effects cellular RNAs.

Observed reductions in cell viability led us to investigate the effect of S411A on infection in mosquitoes. Generally, the longevity of mosquitoes infected with flaviviruses are similar to that of uninfected mosquitoes (45, 46). During mosquito infection, flaviviruses must overcome four barriers: 1) midgut infection barrier, 2) midgut escape barrier, 3) salivary gland infection barrier, and 4) salivary gland escape barrier (47). For the first barrier, the virus must successfully infect and replicate in the midgut epithelial cells (47, 48). Infection is dependent on the arbovirus-specific interactions with the midgut epithelial receptors (49). If the virus cannot establish an infection in the midgut epithelial cells, then the mosquito cannot be infected by the virus. If the virus can establish infection in the midgut, then the next barrier is escaping the midgut by crossing the basal lamina which surrounds the midgut epithelium (47). After escaping the midgut, the virus can disseminate throughout the rest of the mosquito tissues. If the virus is able to penetrate into the salivary gland, the virus must replicate and be deposited into the apical cavities of acinar cells for the mosquito to transmit the virus to other hosts (47). Not all mosquitoes will be able to transmit virus due to unknown reasons. *Culex* mosquitoes in our study were bloodfed or submitted to intrathoracic injection (IT) with either WT or S411A Kunjin. Mosquito mortality was recorded for 15 days for bloodfed mosquitoes or 9 days for IT injected mosquitoes. Results indicated no significant difference in mortality between mosquitoes bloodfed with either WT or S411A Kunjin viruses. Mosquitoes that were intrathoracically injected with S411A Kunjin exhibited an increase in mortality compared to WT Kunjin. Together, our data suggests that S411A Kunjin viruses were inefficient at crossing the midgut infection barrier to establish infection (Fig. 7). However, upon bypassing the midgut infection and midgut escape barriers through IT injection S411A Kunjin was more lethal (Fig. 6B). The basis for the observed increased mortality is not yet clear but could be due to increased cytopathic effect in infected cells similar to what was observed in cell culture.

To further investigate the distribution of WT Kunjin and S411A Kunjin infection within the *Cx. quinquefasciatus* mosquitoes, bodies, legs/wings, and saliva were collected after 7 or 14 days post-bloodfeed and analyzed for the presence of virus. 30 (day 14) to 50% (day 7) of mosquito bodies were positive for WT Kunjin infection, whereas less than 15% of bodies were positive for S411A Kunjin on either collection day. These data suggest that WT Kunjin was able to routinely establish infection within midgut epithelial cells, while S411A Kunjin did so less effectively. However, when legs/wings and saliva were analyzed, WT Kunjin was found at extremely low levels, while S411A Kunjin was found in over half of infected mosquitoes suggesting that once S411A Kunjin was able to cross the midgut escape barrier, it was able to replicate more efficiently in peripheral tissues than WT Kunjin. Previous studies have suggested that arboviruses may require apoptosis to escape the midgut and infect the salivary glands of *Culex* mosquitoes (48, 50–53). Thus, taking into account the cell culture results suggesting S411A Kunjin induces increased cellular death, S411A Kunjin viruses may be able to exit the midgut more effectively than WT Kunjin due to increased induction of apoptosis. Even though S411A Kunjin has a lower initial infection rate, the mutant virus is more toxic to infected cells, and thus, the mutant virus may be able to induce apoptosis and disseminate into the rest of the body leading to a higher potential transmission rate with increased salivary gland infection.

In conclusion, this study provides insight into how a hyperactive NS3 helicase mutant virus contributes to Kunjin virus replication and the effect on cellular responses during infection. S411A Kunjin negatively affects overall replication of the virus and increases the cytopathic effect in cells potentially resulting in increased mosquito mortality. Infection with S411A Kunjin results in less metabolic activity in cells and ultimately cellular death. When considering the increased mortality of mosquitoes IT injected with S411A Kunjin, it seems likely that cells within mosquitoes are undergoing similar cytopathic effect as was observed in cell culture. Cellular death in mosquitoes could allow S411A Kunjin to disseminate into the legs/wings and saliva more efficiently than WT Kunjin and result in increased mosquito death. Virus-induced mortality is not ideal for long-term maintenance of virus in mosquitoes, so flaviviruses appear to have evolved mechanisms to reduce their helicase activity to reduce virus-induced cell killing. Overall, these data indicate that NS3 helicase activity may have significant roles during viral infection in cell culture and *in vivo*, and that NS3 Motif V may play a central role in controlling virus-induced mortality in mosquito vectors to allow for efficient viral transmission.

## MATERIALS AND METHODS

### Cell Culture and Viruses

HEK293T and Vero (African Green Monkey kidney epithelial) cells were maintained in Hyclone Dulbecco’s modified Eagle medium (DMEM) supplemented with 10% fetal bovine serum (FBS), 50 mM HEPES (pH 7.5), 5% penicillin/streptomycin and 5% L-Glutamine. All cells were grown in humidified incubators at 37 °C with 5% CO_2_. The West Nile virus (Kunjin subtype) infectious clone was generously provided from Alexander Khromykh (University of Queensland) (54).

### Virus Mutagenesis

To produce the T407A Kunjin and S411A Kunjin NS3 mutants viruses, a novel bacteria-free virus launch system was used based on *in vitro* NEBuilder assembly of PCR-amplified DNAs containing a eukaryotic Pol II promoter with PCR fragments containing viral genome sequences and direct transfection of assembled DNAs into Vero cells. Three PCR fragments were produced using the Q5 DNA polymerase system (New England Biolabs) according to the manufacturer’s instructions (54). PCR fragment #1 contained the cytomegalovirus (CMV) immediate early promoter (612 bp) using pcDNA-3.1 as the PCR template. PCR fragment #2 (5867 bp) contained the 5’ region of the Kunjin virus genome. PCR fragment #3 (5309 bp) contained the 3’ end of the Kunjin virus genome in addition to a hepatitis delta virus ribozyme. The Kunjin virus infectious clone plasmid FLSDXHDVr was used as the PCR template for fragments #2 and #3 (55). Primer sequences used to produce PCR fragments with overlapping 5’ and 3’ ends for NEBuilder assembly were designed using the NEBuilder Assembly tool (https://nebuilder.neb.com/) and are listed in Table 1.

**Table 1.**
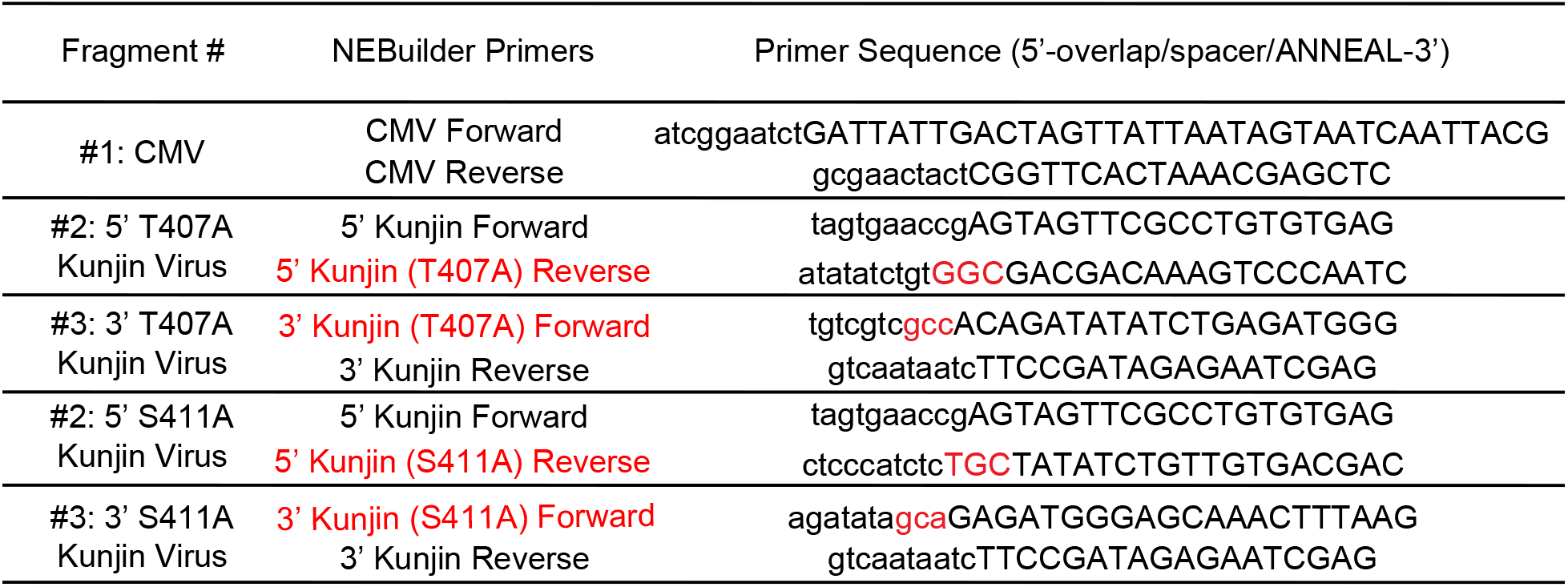
NEBuilder Primers for T407A and S411A Kunjin Viruses. The mutant Kunjin viruses were generated from three fragments: #1, #2, and #3. Primers for fragments #2 and #3 contain the alanine mutation at either position 407 or 411 (highlighted in red). The product of Fragment #2 from the NEBuilder Assembly reaction will contain the specified mutation.

The NS3 T407A and S411A mutations(33) were separately engineered into the Fragment #2 reverse primer and Fragment #3 forward primers. PCR products were gel extracted with the Qiagen Gel Extraction kit and quantified by UV spectrophotometry and agarose gel electrophoresis. To assemble the WT Kunjin, T407A Kunjin, or S411A Kunjin fragments, equal molar amounts of each fragment were mixed in a total DNA mass of 200 ng for each virus in ultrapure water in a final volume of 15 μL. An equal volume of New England Biolabs NEBuilder 2X Master Mix was added to the DNAs, and the reaction was incubated at 50°C for 4 hrs. The assembled DNAs were transfected directly into Vero cells by adding 1 μL of JetPrime transfection reagent (PolyPlus) to the assembly mixture, incubated at 22°C for 15 minutes, and the transfection mixture was added to 50% confluent Vero cells. DMEM media containing 10% fetal bovine serum and 50 mM HEPES (pH 7.5) was changed 24 hours after transfection, and the cells were incubated for 6 additional days and monitored for cytopathic effect. Media was collected on day 6 as the P0 stock. Virus was amplified in a T75 flask seeded at 50% confluency for 7 additional days, and clarified media was collected as the P1 stock. Finally, the P1 stock was used to infect a T150 flask of 50% confluent Vero cells for 7 days, media was collected and clarified of cellular debris, and clarified media frozen at −80°C as the P2 stock. P2 stocks were quantified for infectivity via focus forming assay. T407A Kunjin was unrecoverable from infections. The presence of the S411A Kunjin was verified by extracting RNA from the P2 stock, reverse transcribing and PCR amplifying the NS3 region of Kunjin virus using Kunjin NS3 sequence forward (5’-ATGCACCAATATCCGACTTACA) and reverse (5’-TGGCCTCAGAATCTTCCTTTC) primers, and the sequence of the PCR 794 bp amplicon determine by Sanger sequencing.

### Viral Infectivity

HEK293T cells were plated into 12-well plates at 20,000 cells/well and allowed to adhere to the plates overnight. The next day, the cells were infected at a MOI of 0.01 PFU/cell with either WT Kunjin or S411A Kunjin in triplicate under BSL2 conditions. Both intracellular and extracellular RNA samples were collected every 12 hours for five days. The extracellular RNA samples were processed through focus forming assays to determine the viral titer at each time point. The growth curves were plotting using matplotlib (56).

### Resazurin Assay

HEK293T cells were plated into 96-well plates at 10,000 cells/well. Additionally, DMEM with 10% FBS was plated into one row for each plate as a negative control for resazurin. The following day, cells were either not infected or infected with either WT or S411A Kunjin at a MOI of five PFU/cell. The DMEM media was not infected. Every 24 hours over the course of six days, the cells as well as the negative control were treated with resazurin (0.15 mg/mL). The treated plate was then incubated for 1 hour at 37°C with 5% CO_2_ before measuring the fluorescence at an excitation wavelength of 560 nm and an emission wavelength of 590 nm on a Victor X5 multilabel plate reader (Perkin Elmer).

### CellTiter-Glo Assay

Vero and HEK293T cells were plated into 96-well plates at 10,000 cells/well. The following day, cells in each plate were either not infected or infected with WT or S411A Kunjin at a MOI of five PFU/cell. Every 24 hours for the next six days, cells were treated with 1X of CellTiter-Glo and incubated at room temperature for 10 minutes before measuring luminescence with an exposure time of 0.5 seconds on a Victor X5 multilabel plate reader.

### Mosquitoes

*Cx. quinquefasciatus* mosquito larvae(57), were propagated on a 1:1 mix of powdered Tetra food and powdered rodent chow. Adult mosquitoes were kept on a 16:8 light:dark cycle at 28°C with 70%-80% humidity. Water and sugar were provided ad libitum and citrated sheep blood was provided to maintain the colony. Mosquito infection experiments with Kunjin were performed exclusively on female mosquitoes and under BSL3 conditions.

### Infection of mosquitoes with Kunjin virus and analysis

*Cx. quinquefasciatus* mosquitoes were either fed infectious bloodmeals or intrathoracically injected to introduce Kunjin virus. Bloodfed mosquitoes were fed an infectious bloodmeal of defibrillated calf blood diluted by half with 2.5 × 10^6^ PFU/mL Kunjin virus, or media alone as a negative control. Bloodmeals also contained 2 mM ATP. For IT injection experiments, mosquitoes were injected with 138 nL WT or S411A Kunjin virus (~345 PFU/mosquito) using a Nanoject II (Drummond Scientific). Engorged female mosquitoes were maintained for up to 15 days under conditions described above but in the BSL3 insectary and mortality rate counted daily. For infection, dissemination, and transmission experiments after 7 or 14 days of incubation, mosquitos were cold anesthetized and kept on ice while legs and wings were removed, mosquitoes were salivated for 30 minute in a capillary tube filled with immersion oil, and bodies were collected. Legs/wings and bodies were homogenized at 24Hz for 1 minute in 500 μL mosquito diluent with a stainless steel bead, and saliva samples were stored in 250 μL mosquito diluent as previously described (58). All mosquito samples were clarified by centrifugation at 15,000 X g for 5 minute at 4°C then determined to be positive or negative by infection with undiluted samples by Vero cell plaque assays.

## ACKNOWLEDGEMENTS

We would like to acknowledge the support of NIH grants R01 AI132668 to BJG and R01 AI067380 to GDE. We would also like to acknowledge the helpful discussion with Erin R. Lynch, MS.

